# Inhibition of phosphodiesterase - 10A by Papaverine protects human cortical neurons from quinolinic acid induced oxidative stress and synaptic proteins alterations

**DOI:** 10.1101/2020.12.04.411868

**Authors:** Abid Bhat, Vanessa Tan, Benjamin Heng, Musthafa M. Essa, Saravana B. Chidambaram, Gilles J. Guillemin

**Affiliations:** Department of Pharmacology, JSS College of Pharmacy, JSS Academy of Higher Education & Research, Sri Shivarathreeshwara Nagar, Mysuru, Karnataka 570015, India; Neuroinflammation Group, Faculty of Medicine, Health and Human Sciences, Macquarie University, Sydney, NSW, Australia; Department of Food Science and Nutrition, CAMS, Sultan Qaboos University, Muscat, Oman; Ageing and Dementia Research Group, Sultan Qaboos University, Muscat, Oman; Centre for Experimental Pharmacology and Toxicology, Central Animal Facility, JSS Academy of Higher Education & Research, Mysuru, India

**Keywords:** Phosphodiesterase-10A, cAMP, papaverine, quinolinic acid, oxidative stress, synaptic proteins

## Abstract

Phosphodi esterase-10A (PDE10A) hydrolyse the secondary messengers cGMP and cAMP which play critical role in neurodevelopment and brain functions. PDE10A is linked to progression of neurodegenerative diseases like Alzheimer’s, Parkinson’s, Huntington’s diseases etc and a critical role in cognitive functions. The present study was undertaken to determine the possible neuroprotective effects and the associated mechanism of papaverine (PAP) against quinolinic acid (QUIN) induced excitotoxicity using human primary cortical neurons. Cytotoxicity potential of PAP was analysed using MTS assay. Reactive oxygen species (ROS) and mitochondrial membrane potential were measured by DCF-DA and JC10 staining, respectively. Caspase 3/7 and cAMP levels using ELISA kits. Effect of PAP on the CREB, BNDF and synaptic proteins such as SAP-97, synaptophysin, synapsin-I, PSD-95 expression was analysed by Western blotting technique. Pre-treatment with PAP increased intracellular cAMP and nicotinamide adenine dinucleotide (NAD^+^) levels, restored mitochondrial membrane potential (ΔΨm), and decreased ROS and caspase3/7 content in QUIN exposed neurons. PAP up-regulated CREB and BDNF, and synaptic proteins expression. In summary, these data indicate that PDE10A involves in QUIN mediated neurotoxicity and its inhibition can elicit neuroprotection by reducing the oxidative stress and protecting synaptic proteins via upregulation of cAMP signalling cascade.

## Introduction

Phosphodiestrase-10A (PDE10A) is a key enzyme involved in hydrolysis of intracellular second messengers cyclic adenosine monophosphate (cAMP) and cyclic guanosine monophosphate (cGMP) in brain (Niccolini et al., 2015). cAMP activates protein kinase A (PKA), resulting in the phosphorylation of the transcription factor cAMP response element binding protein (CREB) which in turn induces the protein expression of brain-derived neurotrophic factor (BDNF) and put together these protein regulate a wide range of biological functions, such as synaptic plasticity, learning and memory etc. (Kowiański et al., 2018). Imbalance in cAMP level is implicated in the various neurodegenerative diseases (Roush et al., 2020). Dysregulation of tryptophan metabolism leads to the production of kynurenine metabolites. Quinolinic acid (QUIN) is a neuro- and gliotoxic metabolite produced in kynurenine pathway (KP) (Guillemin, 2012). QUIN causes neurotoxicity by overactivation of N-methyl-D-aspartate (NMDA) receptors and by forming a coordination complex with iron and copper resulting in excessive intracellular Ca^2+^ overload (Chen et al., 2010). It triggers oxidative stress by increasing neuronal nitric oxide synthase (nNOS) and lipid peroxidation (Braidy et al., 2009a). It also impairs mitochondrial oxygen consumption and causes mitochondrial complex (I, II, III and IV) dysfunction (Mishra and Kumar, 2014). Higher levels of QUIN have been found in Alzheimer’s disease (AD) (G. J. Guillemin et al., 2005), Parkinson’s disease (PD) (Zinger et al., 2011), multiple sclerosis (MS) (Lim et al., 2017; Sundaram et al., 2014), Huntington’s disease (HD) (Sumathi et al., 2018). QUIN exposure increases poly (ADP-ribose) polymerase (PARP) activity, depletes NAD^+^ and adenosine triphosphate (ATP) production in human primary neurons (Braidy et al., 2010). Depletion of ATP causes mitochondrial membrane potential (Δψm) collapse in turn the release of cytochrome c (cyt c) and neuronal death (Cao et al., 2011). QUIN intoxication is shown to decreases the synapse number in hippocampus region of the rat via down-regulatingexpression of synaptic proteins such as PSD-95 and reduced phosphorylation of CREB and BNDF expression in rats (Rahman et al., 2018).

Oxidative stress and excitotoxicity suppresses expression of synaptic markers such as synapsin I, synaptophysin, BDNF, calcium□calmodulin dependent protein kinases II and Calcineurin A which hinders brain development and function (Ansari et al., 2008a). PDE10A is an isoenzyme distributed in various brain regions (cerebellum, thalamus, hippocampus, and spinal cord etc); but it is highly expressed in frontal cortex (Heckman et al., 2016). Synaptic density increases in cortical regions after birth in humans (Huttenlocher and Dabholkar, 1997). Interestingly, synaptic proteins are highly expressed in the cortical region and play a vital role in the cortical development (Azir et al., 2018; Valtschanoff et al., 2000). Interaction of pre and postsynaptic proteins facilitate synapse development, communication, long term potentiation and memory formation (Abraham et al., 2019). Alterations in the cortical synaptic proteome is a prominent pathological feature in AD (Counts et al., 2006). Cortical neurons are more vulnerable to ROS attacks which affect synaptic proteins as well (Ansari et al., 2008b). Studies in knockout mice suggests that PDE10A is involved in regulating basal ganglia circuit which governs motor, emotional, and cognitive functions, maintains the energy homeostasis and has a thermoregulatory role (Giampà et al., 2010; Hankir et al., 2016; Siuciak et al., 2006). Its involvment in in neurological disorders such as schizophrenia (Persson et al., 2020) and HD (Giralt et al., 2013) is well documented. Papaverine (PAP), a PDE10A isoenzyme inhibitor, is an major alkaloid obtained from the opium latex of *Papaver somniferum*, family: Papaveraceae (Han et al., 2010) (Fig. 1). PAP is clinically used as a vasodilator and smooth muscle relaxant which mediates its action via cAMP (Kim et al., 2014; Wilson and White, 1986). Phosphodiestrase (PDE) inhibitors are being increasingly considered as therapeutic class for neurological disorders (Bhat et al., 2020). PAP is a potent PDE10A inhibitor with an EC50 value of 36 nM, devoid of any narcotic properties and with lesser side effects as compared to PDE4 inhibitors (Boswell-Smith et al., 2006; Heckman et al., 2016). Understaning the physiological role of PDE10A and advanatges of its inhibitor particualry in brain fucntions, in the present study we investigated the potential effects of PAP against QUIN induced neurotoxicity in human primary cortical neurons. We investigated the effects of PAP on oxidative stress, NAD+/NADH production, Caspase activity, cAMP signaling cascade and synaptic proteins expression in QUIN exposed human neurons.

**Figure 1:**
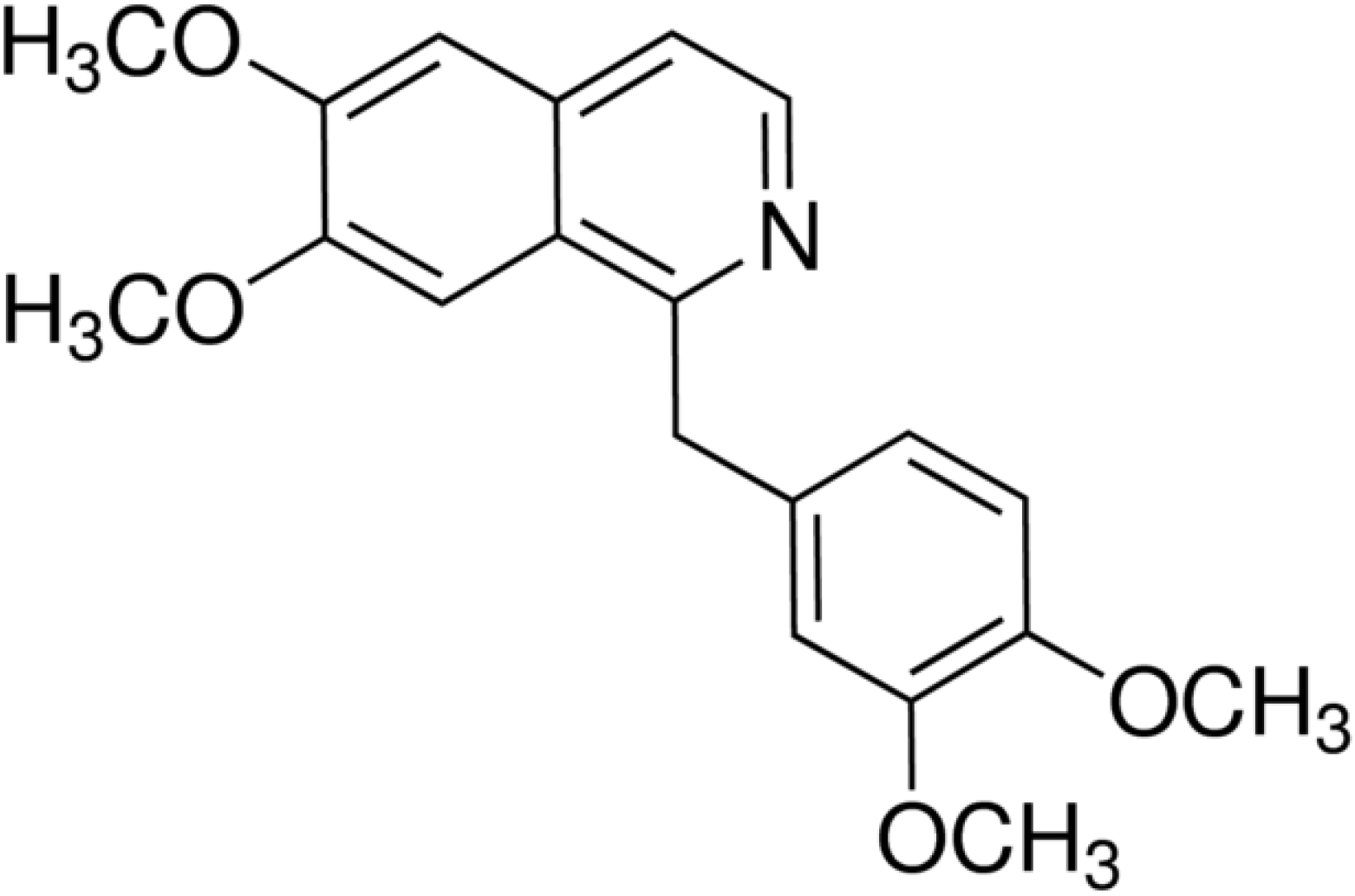
Chemical structure of Papaverine drawn with ChemBioDraw Ultra 12.

## Materials and Methods

### Reagents and antibodies

Papaverine, quinolinic acid, 2’,7’-dichlorofluorescin diacetate (DCFDA) were purchased from Sigma Aldrich (Castle-Hill, Australia). CellTiter 96^®^ Aqueous One Solution cell proliferation assay kit and ApoTox-Glo™ Triplex Assay kit was obtained from Promega, Australia. JC-10 staining kit was obtained from AAT Bioquest, Australia. NAD/NADH Assay Kit, cAMP Elisa kit (ab234585), Rabbit Anti-PSD95 (ab18258), Rabbit Anti-SAP97 (ab3437), Mouse Anti-Synaptophysin (ab8049), Rabbit Anti-BDNF (ab108319) were procured from Abcam, USA. Rabbit Anti-CREB, Rabbit Anti-Synapsin I was obtained from Sigma Aldrich, USA. All other reagents and chemicals used were of analytical grade.

### Primary cortical neuronal culture

Human neuronal cell culture was generated from the 17- to 20-week-old foetal brain tissue collected after therapeutic termination following written informed consent obtained from the participant. Approval for this study was taken from Human Research Ethics Committee of Macquarie University **(Ethics approval: 5201300330).** The cortical neuronal cultures were prepared and maintained according to the method described by Guillemin et al. (2005b). Cells were plated in 96 well and 12-well culture plates coated with Matrigel (1/20 in Neurobasal) and maintained in Neurobasal medium supplemented with 1% B-27 supplement, 1% Glutamax, 1% antibiotic/antifungal (Penicillin G 200 IU/mL, streptomycin sulphate 200 μg/mL), 0.5% HEPES buffer, and 0.5% glucose. The cells were maintained at 37°C in a humidified atmosphere containing 95% air/5% CO_2_ (Guillemin et al., 2007).

### Cell viability assay

Cell viability of papaverine was evaluated using CellTiter 96^®^ AQueous One Solution Reagent (Promega) based on mitochondrial dehydrogenase activity. Neuronal cells were seeded in Matrigel (1/20 in Neurobasal) coated 96 well plate and incubated with a range of PAP concentrations (0.5, 1, 2, 5, 10, 20 μM) at 37°C for 72 hours. Twenty microliters of [3-(4,5-dimethylthiazol-2-yl)-5-(3-carboxymethoxyphenyl)-2-(4-sulfophenyl)-2H-tetrazolium (MTS) solution was added to each well. Absorbance was read at 490 nm in a microplate reader (PHERAstar FS) at 24, 48 and 72 hours after PAP exposure.

### Treatment protocol

Human primary neurons were pre-treated for 24 h with papaverine (2 and 5 μM; doses were fixed based on cytotoxicity assay results) followed by 48 hours exposure with QUIN (2 μM) at 37°C.

### Reactive Oxygen Species detection using 2-7-Dichlorofluorescin Diacetate Assay

Intracellular reactive oxygen species levels in primary human neurons was estimated using 2-,7-dichlorofluorescin (DCF)-DA assay. (Kim et al., 2012). After completing the treatment, cells were washed with ice cold phosphate-buffered saline (PBS). DCF-DA (20 μM) was added and incubated for 30 minutes. Fluorescence intensity was measured using a microplate reader (PHERAstar FS, BMG Labtech) (excitation wavelength of 495 nm and emission wavelength of 515 nm). ROS production was calculated as a percentage of the control.

### Mitochondrial membrane potential (ΔΨm) assessment

Mitochondrial membrane potential was determined by using highly sensitive JC-10 (tetraethyl benzimidazolylcarbocyanide iodine) staining as per the manufacturer’s instructions (Li et al., 2016). The accumulation of the JC-10 dye is proportional to the mitochondrial membrane potential. JC-10 gives green and red to orange fluorescence in low and high ΔΨ*m*, respectively. Following the pre-treatment with PAP, cells were washed with ice cold PBS. 100 μL of JC-10 stain (30 μM) added and incubated for 20 minutes. JC-10 staining in the culture media was removed and 100 μL of HEPES buffer was added. The change in fluorescence intensity was measured using microplate reader (PHERAstar FS) at Ex/Em = 490/525 nm and 540/590nm. The shift from green to red indicates depolarisation.

### Measurement of caspase 3/7 activity

Caspase 3/7 activity in primary neuronal cells was measured using ApoTox-Glo Triplex Assay kit (Promega, Madison, WI, USA) as per the manufacture’s instruction. Cells were treated with PAP and QUIN as mentioned in the treatment protocol. 50 μL of caspase Glo 3/7 reagent was added to each well and incubated for 2 hours at room temperature with constant shaking. Caspase 3/7 activity was analysed by measuring luminescence using a microplate reader (PHERAstar FS).

### Estimation of intracellular NAD^+^/NADH levels

Intracellular NAD^+^/NADH ratio were measured using NAD/NADH Assay Kit (Abcam, Inc., Cambridge, MA, USA) as per the instructions (Ren et al., 2010, p. 2). Cells were washed with ice cold PBS following the completion of treatment protocol. The cells were lysed with 400 μL of NAD^+^/NADH extraction buffer containing a cocktail of protease inhibitor (Roche Diagnostic, Castle Hill, NSW, Australia) and centrifuged at 10000×g for 5 min at 4°C. To measure NAD^+^, 200 μL aliquot of cell lysate was heat quenched at 60°C for 30 min. An equal volume of NAD^+^/NADH reaction mixture was added to each well with cell lysate, incubated at room temperature for 60 min, and absorbance was read at 450 nm using a microplate reader (PHERAstar FS). The concentration of NAD^+^ or NADH was calculated using a standard calibration curve.

### Measurement of cAMP concentration

Cells were seeded in 12 well plate and were pre-treated with vehicle or PAP (2 and 5 μM) for 24 hours and then exposed to QUIN (2 μM) for 48 hours. At the end of the treatment protocol, cells were washed with ice cold PBS and lysed using 0.1 M HCl for 20 minutes and centrifuged for 10 minutes. Cell lysate was collected, and cAMP concentration was measured using a direct immunoassay kit (Abcam, MA) following the manufacturer’s instruction. In brief, standards and samples were added to wells coated with an IgG antibody. A cAMP conjugated to alkaline phosphatase was then added, followed by a rabbit polyclonal antibody against cAMP. The antibody binds to cAMP in the sample or to the conjugate in a competitive manner. The plate was washed, leaving only bound cAMP. para-Nitrophenyl phosphate (pNpp) substrate solution was added and produced a yellow color when catalyzed by the alkaline phosphatase on the cAMP conjugate. The stop solution was then added, and the yellow color was read at 405 nm using a microplate reader (PHERAstar FS), and concentration of cAMP was calculated using a standard calibration curve.

### Western blot analysis

Cells were lysed with radioimmunoprecipitation assay (RIPA) buffer (50 mM Tris, pH 7.4, 150 mM NaCl, 1% NP-40, 5 mM EDTA, 0.5% sodium deoxycholate, 0.1% SDS, 50 nMNaF, 1 mM sodium vanadate) containing a cocktail of protease inhibitor (Roche Diagnostic, Castle Hill, NSW, Australia). Total protein concentration was determined by BCA protein assay (Bio-Rad Laboratories, Hercules,CA, USA), cell lysate samples were aliquoted and stored at −80°C till used. Proteins (20 μg) were separated by using 12% bis-tris −SDS-PAGE (NuPAGE, Invitrogen, Carlsbad CA, USA) by electrophoresis. Resolved proteins in the gels were transferred onto nitrocellulose membranes (Biorad) and electroblotted. Membranes were blocked with 5% non-fat skimmed milk in Tris-Buffered Saline and Tween 20 (TBST) for 1 hour followed by overnight incubation with the primary antibodies CREB (1:1000), BDNF (1:1000), PSD-95 (1:1000), Synapsin (1:1000), Synaptophysin (1:1000), SAP 97 (1:1000) at 4°C. The membranes were rinsed with TBST (3 washings for 10 minutes), incubated with the secondary antibodies (HRP conjugated anti-mouse or anti-rabbit IgG) for 1 h at room temperature followed by 3 washings for 10 mins with TBST. The bands were visualised by Odyssey Infrared Imaging System; Li-Cor, Lincoln, NE

### Statistical analysis

Data were presented as mean ±SEM. Group mean differences were analysed using one-way ANOVA test followed by Tukey’s multiple comparison test as post hoc test. Results were analysed using GraphPad Prism version 7.04 with probability value of *p* ≤0.05 considered as significant.

## Results

### Cell viability

Cell viability was tested using CellTiter 96^®^ AQueous One Solution Reagent (Promega) after 24-, 48 and 72-hour treatment with papaverine at concentrations 0.5, 1, 2, 5, 10 and 20 μM. **(Fig.2).** We did not observe time or concentration dependent neuronal death in cell viability assay

**Figure 2:**
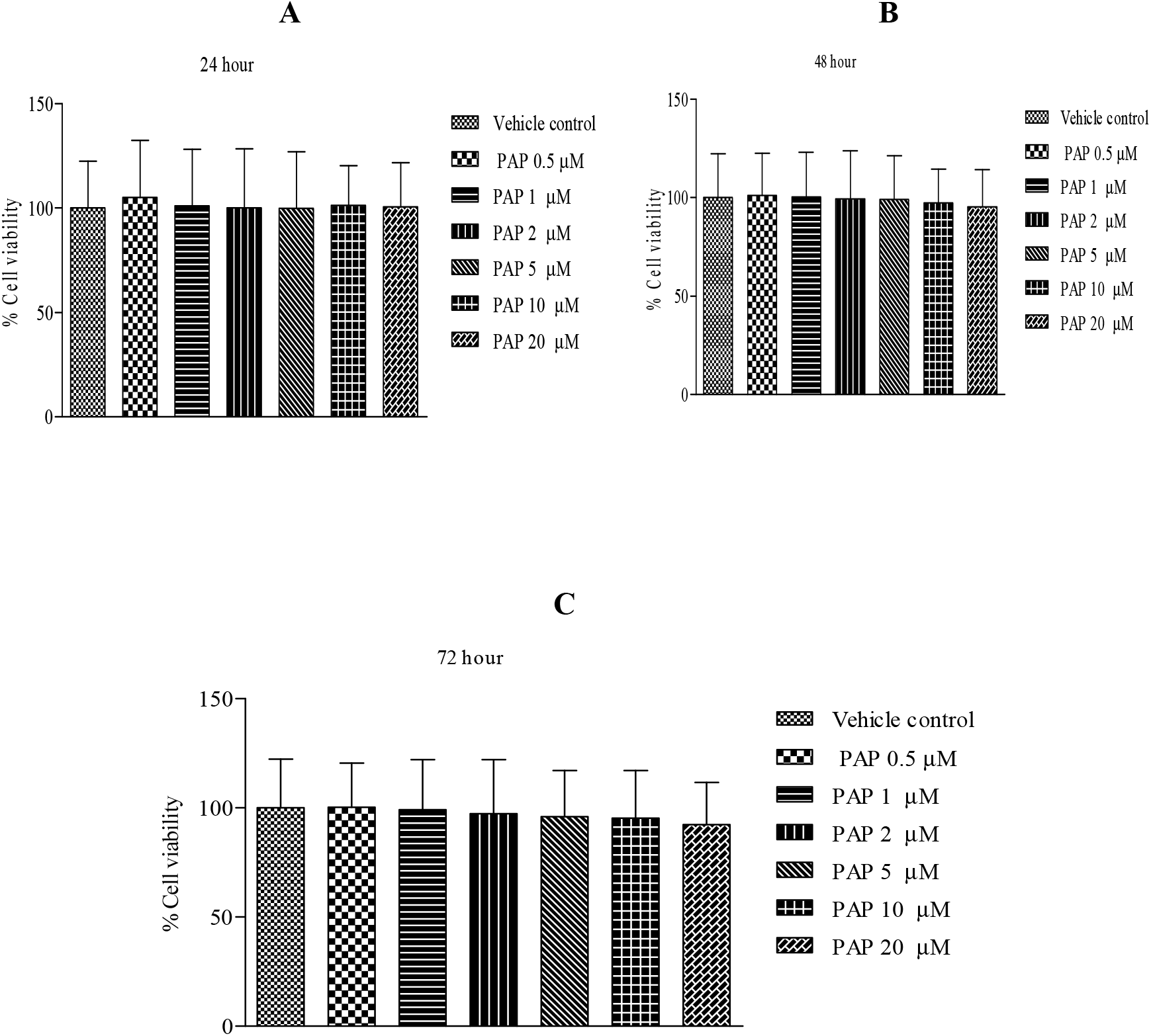
Papaverine did not show toxic effects at the tested concentration in human cortical neurons. (A) Human cortical neurons were treated with various concentrations of PAP and assessed for cell viability at 24 (A), 48 (B) and 72 h (C) using MTS solution.

### Papaverine reduces QUIN induced reactive oxygen species (ROS) generation in human cortical neurons

QUIN exposed neurons showed significant (*p* <0.01) increase in ROS production compared to vehicle treated cortical neurons. Pre-treatment with PAP reduced ROS production in QUIN intoxicated neurons. But a significant (*p* <0.01) decrease in ROS production was observed at 5 μM concentration **(Fig. 3)**. This shows that PAP has the potential to reduce oxidative stress.

**Figure 3:**
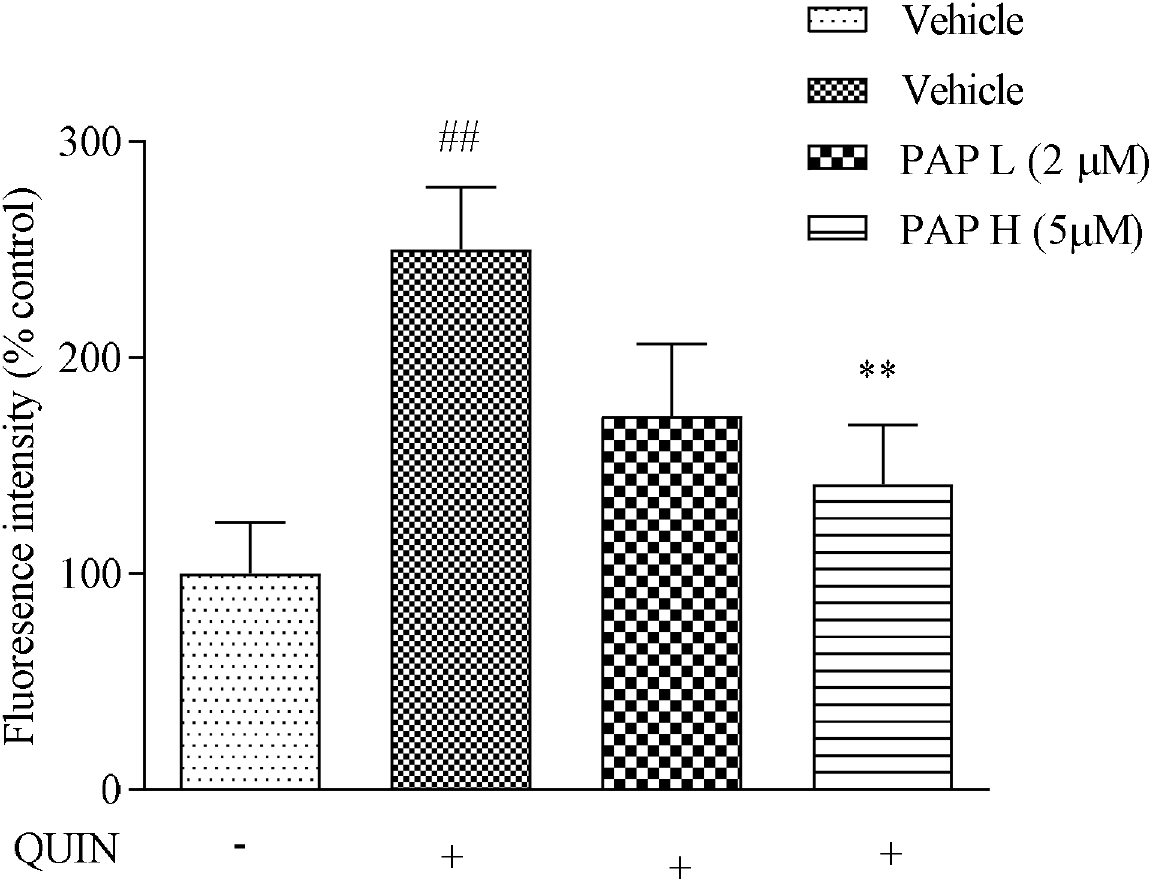
Papaverine reduces QUIN-induced ROS production in human neurons. Neurons were pre-treated with PAP (2μM & 5 μM) for 24 hours, followed by 48-hour exposure with QUIN (2μM). Data are presented as mean ± SEM (n = 3) and represent three independent experiments. ## denotes *p*< 0.01 vs vehicle, ** denotes *p*< 0.01 vs QUIN exposed neurons

### Papaverine restores mitochondrial membrane potential (ΔΨm) in QUIN exposed human neurons

Mitochondrial oxidative stress and ΔΨm are involved in the neurodegeneration. Amelioration of mitochondrial dysfunction is proposed as an effective way for slowing down the progression of neurodegeneration (Wu et al., 2019). Increased production of ROS affects mitochondrial structure and function, we assessed ΔΨm by using JC10 staining. Mitochondrial depolarization results in dye release and unquenching, increasing the fluorescence signal is proportional to ΔΨm values. In QUIN exposed human neurons we found that fluorescence intensity increased significantly (*p* < 0.01) when compared with vehicle treated neurons, indicating increased mitochondrial depolarisation. Pre-treatment with PAP decreased mitochondrial depolarization in QUIN intoxicated neurons with a significant (p <0.05) protection recorded at 5μM **(Fig. 4).**

**Figure 4:**
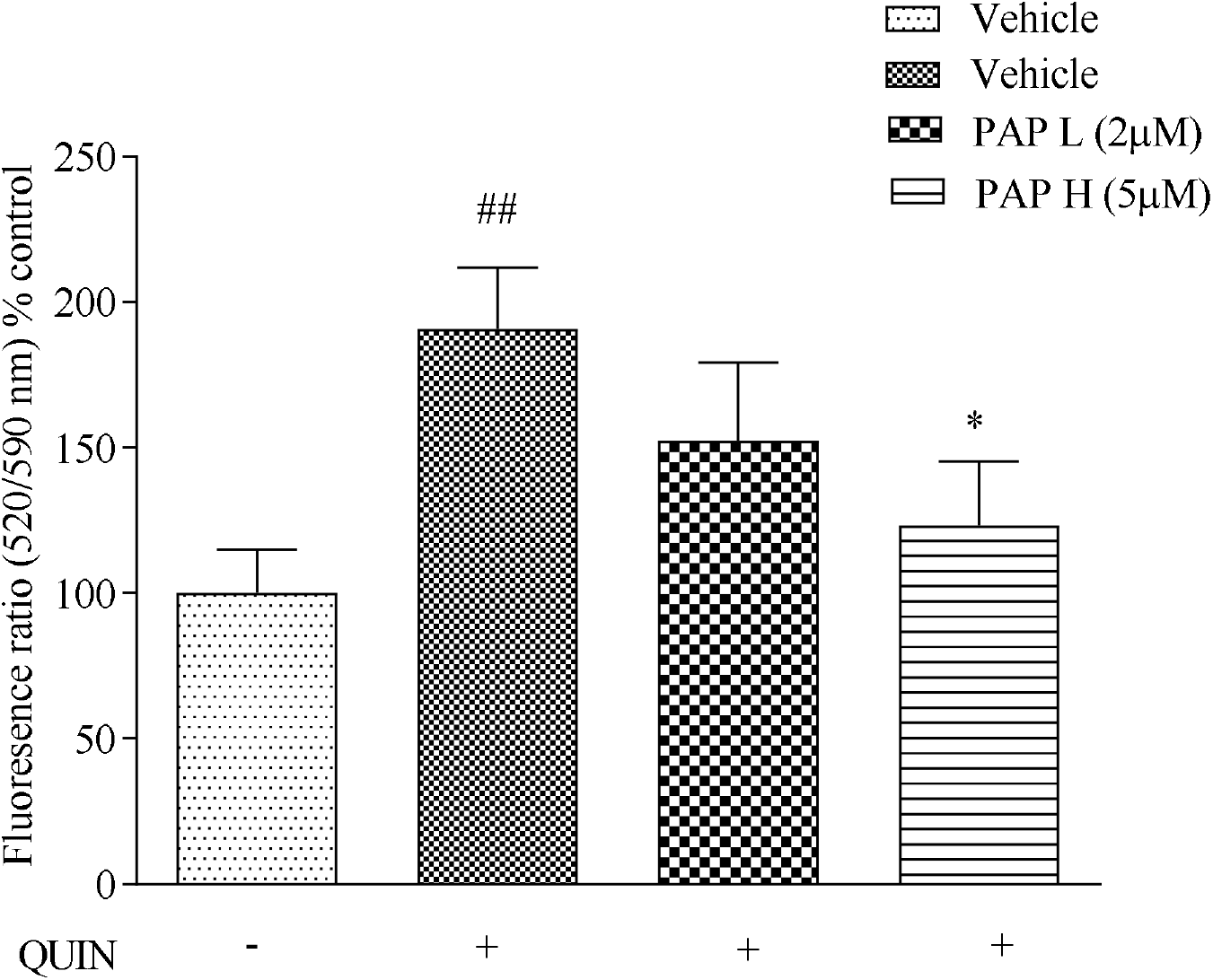
Papaverine restores mitochondrial membrane potential in QUIN exposed human cortical neurons. Neurons were pretreated with PAP (2μM & 5 μM) for 24 hours, followed by 48-hour exposure with QUIN (2μM). Neurons were incubated with JC-10 (30 μM) for 20 min. Data are presented as mean ± SEM (n = 3) and represent three independent experiments. ## denotes *p*< 0.01 vs vehicle, * denotes *p*< 0.05 vs QUIN exposed neurons

### Papaverine suppressed QUIN induced caspase 3/7 activity in human neurons

Neurons primarily use the intrinsic apoptotic pathway to undergo cell death and role of caspase-3 and caspase 7 in progression of neurodegeneration is well studied using knockout mouse model (D’Amelio et al., 2010). Increase in the oxidative stress elicits neurodegeneration by activating the early apoptosis cascade (Dos Santos et al., 2018). We found that QUIN significantly (p < 0.01) increased caspase3/7 activity in neurons when compared to vehicle treated cells. Pre-treatment with papaverine reduced QUIN induced caspase 3/7 activity with a significant (*p* < 0.01) decrease at 5 μM concentration when compared with QUIN alone exposed cells (**Fig.5).**

**Figure 5:**
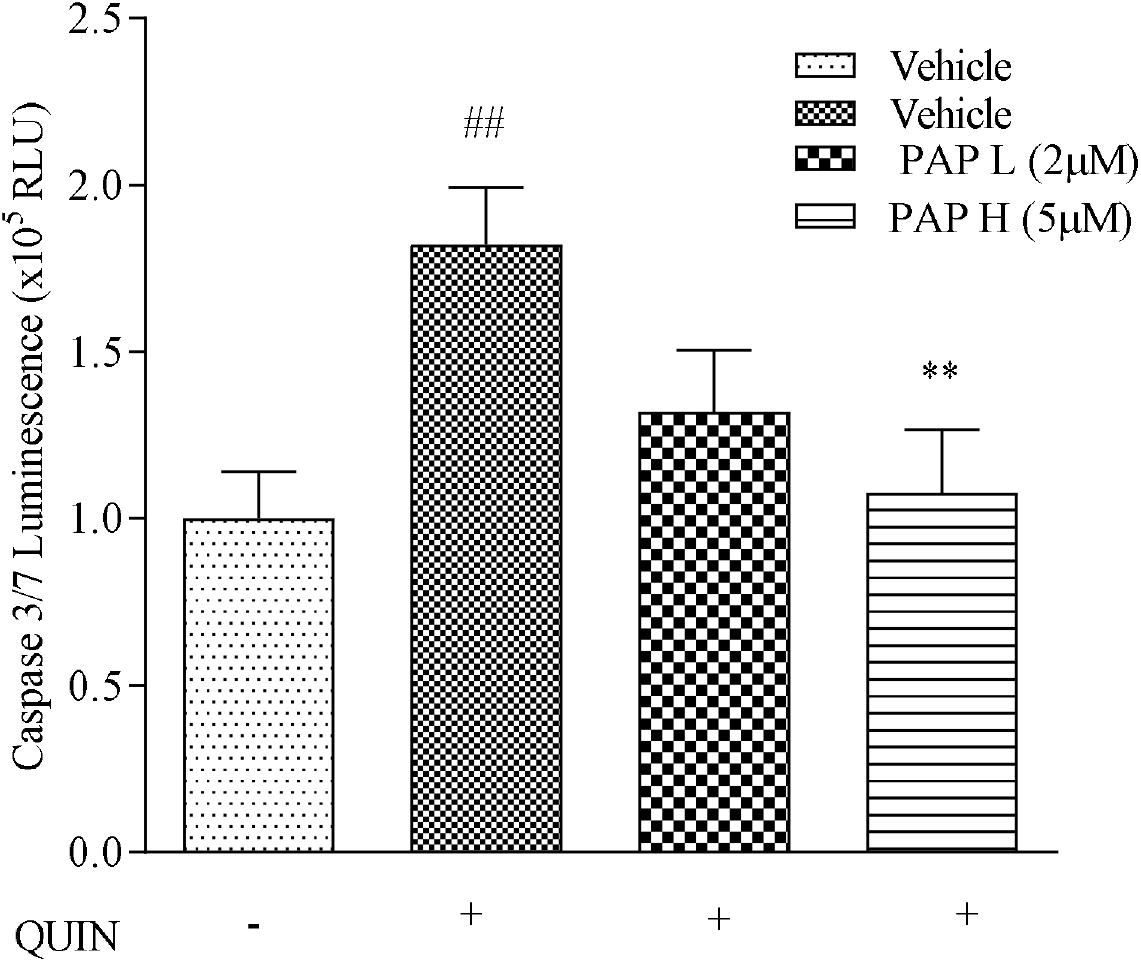
Papaverine decreases Caspase 3/7 activity in QUIN exposed human neurons. Neurons were pre-treated with PAP (2μM & 5 μM) for 24 hours, followed by 48-hour exposure with QUIN (2μM). Caspase 3/7 activity was assessed using ApoTox-Glo Triplex Assay kit (Promega, Madison, WI, USA). Data are presented as mean ± SEM (n = 3) and represent three independent experiments. ## denotes *p*< 0.01 vs vehicle, **denotes *p*< 0.01 vs QUIN exposed neurons

### Papaverine increases intracellular NAD^+^/NADH level in human neurons

We also investigated the effects of PAP on NAD/NADH content in QUIN treated cells. Human neurons exposed to QUIN showed significant (*p* < 0.01) decrease in intracellular NAD^+^/NADH content when compared with the vehicle treated cells. Papaverine pre-treatment showed dose dependent increase in NAD^+^/NADH content and a significant (*p* < 0.01) increase was found at 5 μM when compared with QUIN exposed human neurons **(Fig. 6).**

**Figure 6:**
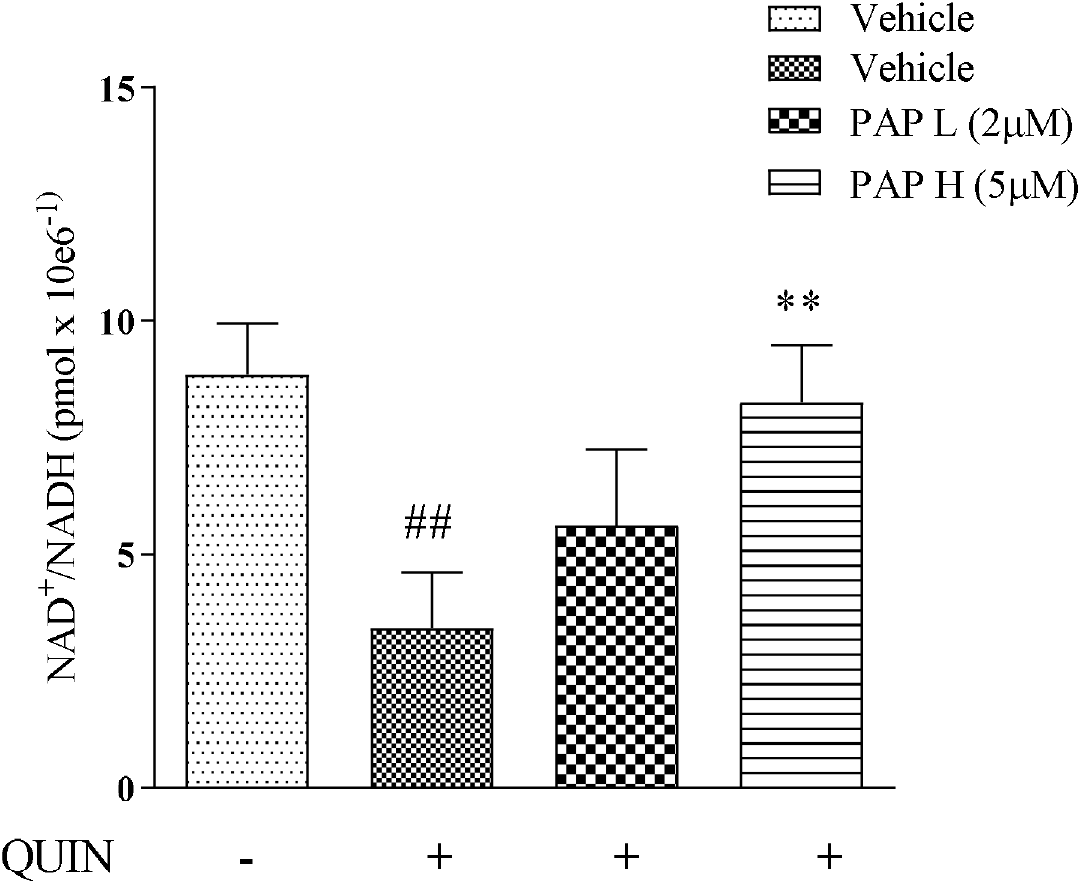
Papaverine increases NAD^+^/NADH levels in QUIN exposed human neurons. Neurons were pre-treated with PAP (2 μM & 5 μM) for 24 hours, followed by 48-hour exposure with QUIN (2μM). NAD^+^/NADH concentration were measured in accordance with the manufacturer’s instruction (Abcam, Inc., Cambridge, MA, USA). Data are presented as mean ± SEM (n = 3) and represent three independent experiments. ## denotes *p*< 0.01 vs vehicle, ** denotes *p*< 0.01 vs QUIN exposed neurons

**Figure 6:**
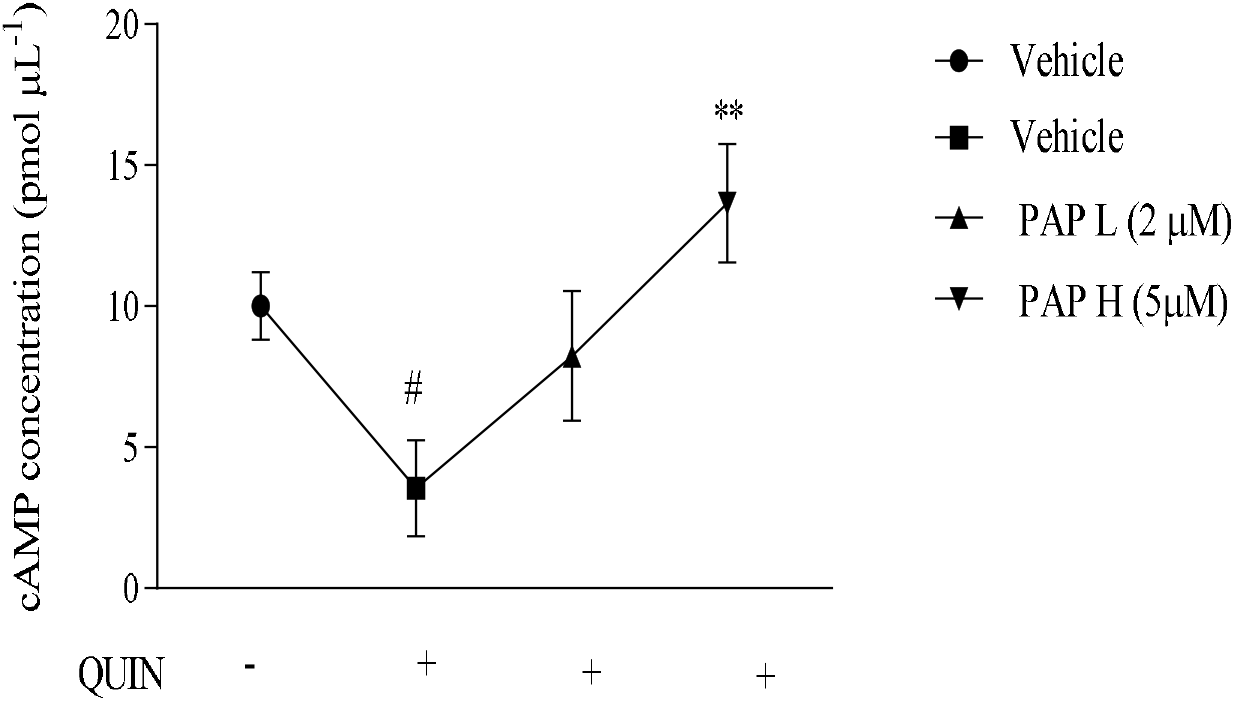
Effect of PAP on cAMP concentration in QUIN exposed human neurons. Pre-treatment with PAP increased cAMP concentration in QUIN exposed human neurons. Data are presented as mean ± SEM (n = 3) and represent three independent experiments. #denotes *p*< 0.05 vs vehicle, **denotes *p*< 0.01 vs QUIN exposed neurons

### Papaverine increases intracellular cAMP content in QUIN exposed human neurons

Next, we investigated the effects of PAP on cAMP content in human neurons using ELISA kit. QUIN exposed neurons were found to have a significant (*p* < 0.05) decrease in cAMP levels as compared to vehicle treated neurons. Pre-treatment with PAP increased the cAMP levels when compared with QUIN exposed neurons with a significant (*p* < 0.01) increase at 5 μM concentration **(Fig.7).**

**Figure 7:**
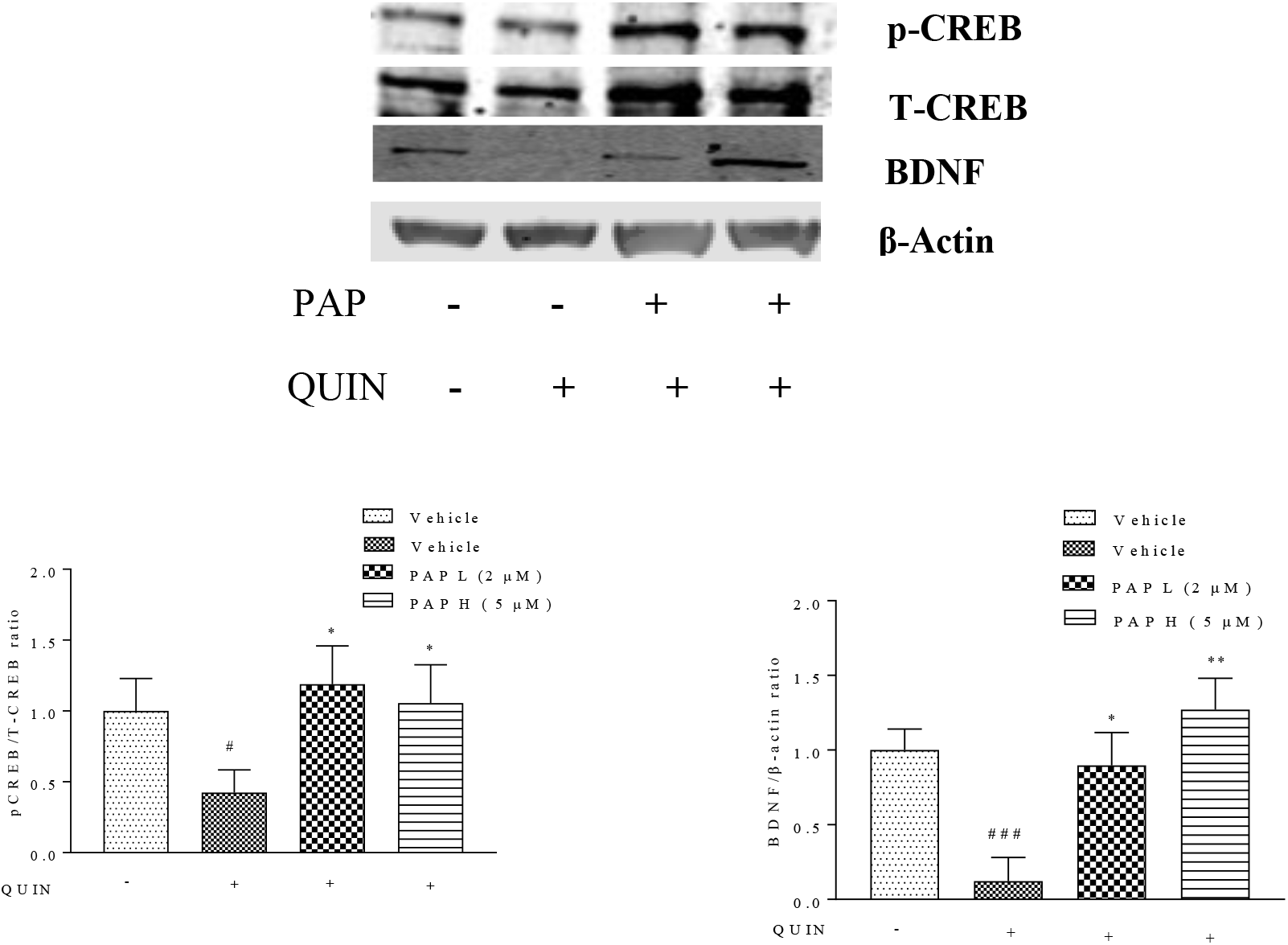
PAP upregulates the expression of CREB and BDNF in QUIN exposed neurons. Neurons were pre-treated with PAP (2μM & 5 μM) for 24 hours, followed by 48-hour exposure with QUIN (2μM). (A) Quantification of pCREB/T-CREB. (B) Quantification of BDNF/ β-actin. Data are presented as mean ± SEM (n = 3) and represent three independent experiments. # denotes *p*< 0.05, ###denotes *p*< 0.001 vs vehicle, * denotes *p*< 0.05, ** denotes *p*< 0.01 vs QUIN exposed neurons

### Papaverine upregulated CREB/ BDNF expression in primary human neurons

Further we investigated the effects of PAP on the expression of CREB and BDNF in QUIN exposed human cortical neurons. CREB is an important cellular transcription factor regulating the expression of neurotrophic factors such as BDNF are important for synaptic plasticity and memory (Wang et al., 2018). QUIN was shown to significantly (*p* < 0.05) reduce the expression of CREB in human neurons. Pre-treatment with papaverine significantly increased (*p* <0.05) the expression of CREB in human neurons **(Fig.8A).** Brain-derived neurotrophic factor (BDNF) is reported to play an important role in the survival of neurons (Miranda et al., 2019). A significant decrease (*p* < 0.05) in BDNF expression was found in QUIN exposed human neurons. Pre-treatment with papaverine significantly (*p* < 0.01) reversed the QUIN induced decline in BDNF expression in human neurons as compared to QUIN exposed cells **(Fig.8B)**.

### Papaverine increased the expression of synaptic associated proteins like Synapsin-I, Synaptophysin, PSD-95 and SAP-97

Next, we investigated the effects of PAP against QUIN mediated synaptic damage in human neurons. QUIN exposure significantly reduced the expression of Synapsin I (*p < 0.05*), synaptophysin (*p < 0.01*), PSD-95 (*p < 0.01*) and SAP-97 (*p < 0.05*) in human neurons. Pre-treatment with PAP significantly upregulated the expression of Synapsin I (*p < 0.01*) synaptophysin (*p < 0.01*), PSD-95 (*p < 0.01*) and SAP-97 (*p < 0.01*) when compared with QUIN exposed neurons **(Fig. 9).** Taken together, these results indicate that inhibition of PDE10A with papaverine was enough to induce both presynaptic and postsynaptic remodelling and instigate the expression of synapsis associated proteins in QUIN exposed human neurons.

**Figure 9:**
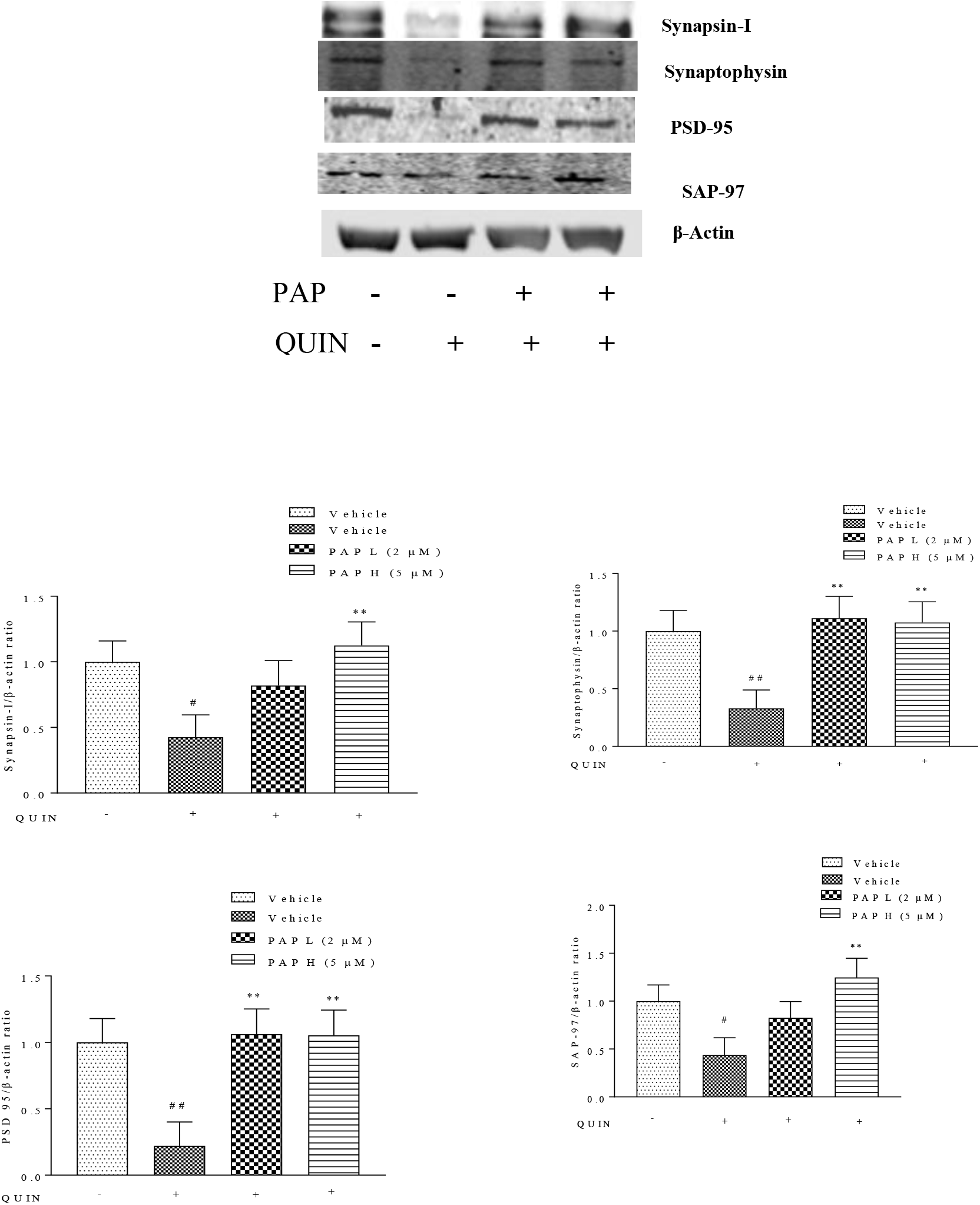
Papaverine increased the expression of Presynaptic and postsynaptic proteins in QUIN exposed neurons. Neurons were pre-treated with PAP (2μM & 5 μM) for 24 hours, followed by 48-hour exposure with QUIN (2μM). (A) Quantification of Synapsin-I-97/β-actin. (B) Quantification of Synaptophysin/ β-actin (C) Quantification of PSD-95/ β-actin (D) Quantification of SAP-97/ β-actin. Data are presented as mean ± SEM (n = 3) and represent three independent experiments. # denotes *p*< 0.05, ##denotes *p*< 0.01 vs vehicle, ** denotes *p*< 0.01 vs QUIN exposed neurons

## Discussion

The present study is the first of its kind to show the protective effects of papaverine against QUIN induced excitotoxicity in human neurons. Our results showed that PAP alleviates oxidative stress by upregulating cAMP and enhances the expression of synaptic proteins. PDE10A is highly expressed in the hippocampus and cortex which are highly vulnerable to excitotoxicity (Heckman et al., 2016). PDE10A plays a pivotal role in these neurons as it regulate cAMP cascade and its inhibition is shown to activate and phosphorylate CREB and other neurotrophic factors (Steffan et al., 2000). Increased expression of PDE10A precipitates NMDA receptors and increased dopamine D_1_- and D_2_ receptor activity which alters cognitive and motor functions by disrupting the integration of information from cortical projections (Smith et al., 2013). Several studies have shown that PDE10A is implicated in the progression of neurodegenerative diseases such as HD (Cardinale and Fusco, 2018; Harada et al., 2017), PD (García et al., 2014; Lee et al., 2019a), MS (Suzumura et al., 1999) schizophrenia (Schmidt et al., 2008) and cognitive dysfunctions (Rodefer et al., 2012) which is due to overexcitation of NMDA receptors and disruption of mitochondrial bioenergetics (Bading, 2017).

Oxidative stress causes destruction of genetic materials, lipids and proteins (Birben et al., 2012) and also linked to the pathogenesis of many neurodegenerative diseases (Islam, 2017). Recenty we compiled the data that sleep deprivation alters tryptophan metabolism and the neurotoxic KP metabolites produced impairs the cognitive functions (Abid, 2020). In this study we used QUIN to induce oxidative stress in human neurons and observed that PAP, a cAMP specific PDE10A inhibitor, has promising neuroprotective effects via improving cAMP signalling and synaptic proteins expression. QUIN increases PARP activity, depletes NAD^+^ (Braidy et al., 2009b), ATP production and activates apoptotic cascade resulting in neuronal death (Gilles J Guillemin et al., 2005). Inhibition of PDE class of enzymes have been found to reduce the apoptosis in neuronal cells (Mizuno et al., 2004). In the present study QUIN exposure induced ROS production, caused mitochondrial membrane depolarization, increased Caspase3/7 activity, and reduced NAD^+^/NADH content in human primary neurons. Papaverine administration have been found to reduced Caspase-3 expression (Yurtsever Kum et al., 2018) and lipid peroxidation in rats (Chandra et al., 2000). Papaverine is reported to restore mitochondrial respiration (Benej et al., 2018) by inhibiting ROS production via supressing the expression of p47phox and improving Nrf2/ARE signalling cascade in PD mouse model (Lee et al., 2019b). Consistently, in the present study, we also observed a decrease in the production of ROS, restoration of mitochondrial membrane potential, decrease in caspase3/7 activity and increased production of NAD^+^ in human neurons treated with papaverine, which reveals the crucial role of PDE10A on oxidative stress and apoptosis. cAMP signalling cascade contributes in reducing inflammation and oxidative stress (Jung et al., 2010). Upregulation of PKA/CREB is closely linked with the ROS neutralising, reducing neuroinflammation and increasing mitochondrial biogenesis (Fernandez-Marcos and Auwerx, 2011). PAP improved cognitive function in HD mice by upregulating the expression of cAMP/CREB in hippocampal region (Giralt et al., 2013). It is reported to increase the expression of neurotrophic factors like BDNF and GNDF which are necessary for the neuronal survival (Lee et al., 2019b). Increased expression of BDNF facilitates long term potentiation and enhances memory consolidation in mice (Radiske et al., 2017). Earlier we showed that PDE4 inhibition upregulates CREB/BDNF expression in renovascular hypertensive rats (Jabaris et al., 2015). Similarly, PDE10A inhibition with PAP shown to exert neuroprotective effect by supressing microglial activation via nuclear factor kappa-light-chain-enhancer of activated B cells (NF-κB) signaling pathway and proinflammatory mediators and upregulates peroxisome proliferator-activated receptor gamma (PPARγ) signaling in mice (Dang et al., 2016; Lee et al., 2019b).

The current study adds evidence that PDE10A inhibition also increases cAMP/CREB and BDNF expression in the QUIN intoxicated human neurons. Further, cAMP is shown to influences the expression of synaptic proteins which are necessary for synaptic transmission and release of neurotransmitters (Leenders and Sheng, 2005). Activation of cAMP/PKA/CREB cascade enhances synaptic transmission in hippocampal neurons (Leenders and Sheng, 2005). Furthermore, BDNF upregulates the expression of presynaptic and postsynaptic proteins such as PSD-95, SAP-97, synaptophysin in cortical neurons of Sprague-Dawley rats (Jourdi and Kabbaj, 2013). We have also recorded a significant increase in the expression of presynaptic and postsynaptic proteins like SAP-97, synaptophysin, synapsin-I and PSD-95 with PAP pre-treatment in QUIN exposed human cortical neurons. Thus it can be inferred that PAP produces neuroprotection by upregulating cAMP signalling cascade which combats the oxidative stress and synaptic dysfunction. Further in vivo studies are in progress in our lab to study the effects of papaverine on synaptic proteins expression and cognition in sleep deprived mice, wherein altered kynurenine metabolism and increased QUIN level produces neurotoxicity.

## Conclusion

The present study reports the neuroprotective effects of papaverine, a PDE10A inhibitor, against quinolinic acid induced excitotoxicity in human neurons. Our study suggests that upregulation of cAMP cascade by papaverine plays a key role in reducing oxidative stress and increasing the expression of synaptic proteins. Therefore, papaverine may be considered a promising therapeutic candidate for further studies aiming to improve synaptic function in neurodegenerative diseases.

## Acknowledgment

Mr. Abid Bhat is supported by the IBRO-APRC Exchange Fellowships program, Macquarie University and Senior Research Fellowship from the Indian Council of Medical Research (ICMR, New Delhi, India). Prof Guillemin is supported by the National Health Medical Research Council (NHMRC), the Australian Research Council (ARC), the Hanbury Foundation, the Mason Foundation and Macquarie University.

## Informed Consent

Written informed consent was obtained from the parents (5201300330).

## Abbreviations

AD: Alzheimer’s diseases
ALS: Amyotrophic lateral sclerosis
ANOVA: Analysis of variance
Aβ: Amyloid beta
ARE: Antioxidant response element
ATP: Adenosine triphosphate
BDNF: Brain-derived neurotrophic factor
cAMP: Cyclic adenosine monophosphate
cGMP: Cyclic guanosine monophosphate
CREB: cAMP response element-binding protein
Cyt c: Cytochrome c
DCFDA: 2□, 7□-Dichlorofluorescin Diacetate
EDTA: Ethylenediaminetetraacetic acid
HEPES: 4-(2-hydroxyethyl)-1-piperazineethanesulfonic acid
HD: Huntington’s disease
JC-10: 5,5,6,6’-tetrachloro-1,1’,3,3’ tetraethylbenzimidazoylcarbocyanine iodide
KP: Kynurenine pathway
MS: Multiple sclerosis
MTS: 3-(4,5-dimethylthiazol-2-yl)-2,5-diphenyltetrazolium bromide
NAD: Nicotinamide adenine dinucleotide
NMDA: N-methyl-D-aspartate
NF-Kb: Nuclear factor kappa-light-chain-enhancer of activated B cells
nNOS: Neuronal nitric oxide synthase
Nrf2: Nuclear erythroid 2-related factor 2
PAP: Papaverine
PARP: Poly (ADP-ribose) polymerase
PBS: Phosphate-buffered saline
PD: Parkinson’s disease
PDE: Phosphodiesterase
PDE10A: Phosphodiesterase-10A
PKA: Protein kinase A
pNpp: para-Nitrophenyl phosphate
PPARy: Peroxisome proliferator-activated receptor gamma
PSD-95: Post synaptic density protein-95
QUIN: Quinolinic acid
RIPA: Radioimmunoprecipitation assay
ROF: Roflumilast
ROS: Reactive oxygen species
SAP 97: Synapse-associated protein 97
SDS: Sodium dodecyl sulphate
SYN1: Synapsin-I
TBST: Tris-Buffered Saline and Tween 20
ΔΨm: mitochondrial membrane potential

---

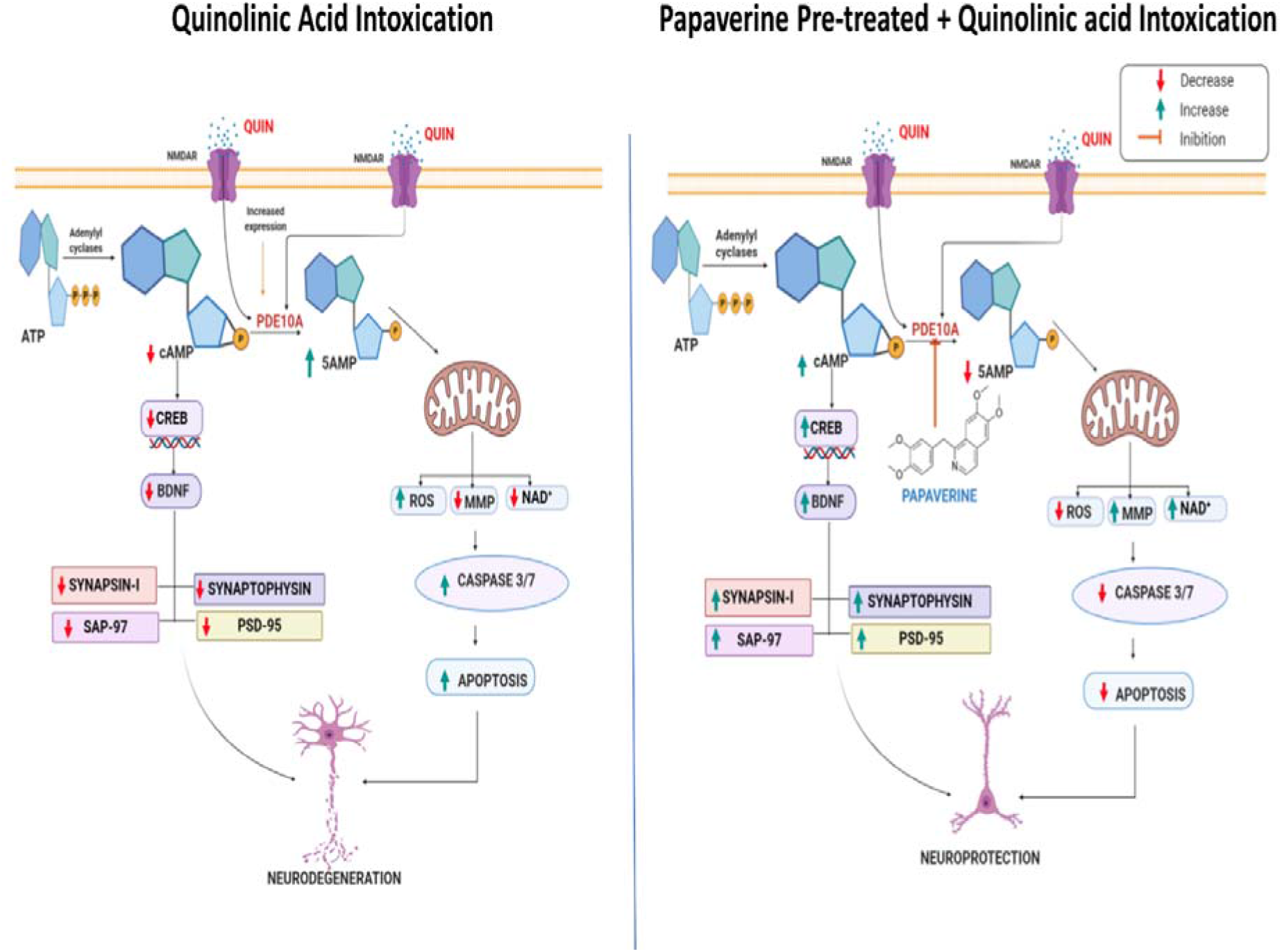

## References

Abraham, W.C., Jones, O.D., Glanzman, D.L., 2019. Is plasticity of synapses the mechanism of long-term memory storage? Npj Sci. Learn. 4, 1–10. https://doi.org/10.1038/s41539-019-0048-y

Ansari, M.A., Roberts, K.N., Scheff, S.W., 2008a. A time course of contusion-induced oxidative stress and synaptic proteins in cortex in a rat model of TBI. J. Neurotrauma 25, 513–526. https://doi.org/10.1089/neu.2007.0451

Ansari, M.A., Roberts, K.N., Scheff, S.W., 2008b. Oxidative stress and modification of synaptic proteins in hippocampus after traumatic brain injury. Free Radic. Biol. Med. 45, 443–452. https://doi.org/10.1016/j.freeradbiomed.2008.04.038

Bading, H., 2017. Therapeutic targeting of the pathological triad of extrasynaptic NMDA receptor signaling in neurodegenerations. J. Exp. Med. 214, 569–578. https://doi.org/10.1084/jem.20161673

Benej, M., Hong, X., Vibhute, S., Scott, S., Wu, J., Graves, E., Le, Q.-T., Koong, A.C., Giaccia, A.J., Yu, B., Chen, C.-S., Papandreou, I., Denko, N.C., 2018. Papaverine and its derivatives radiosensitize solid tumors by inhibiting mitochondrial metabolism. Proc. Natl. Acad. Sci. U. S. A. 115, 10756–10761. https://doi.org/10.1073/pnas.1808945115

Bhat, A., Ray, B., Mahalakshmi, A.M., Tuladhar, S., Nandakumar, D., Srinivasan, M., Essa, M.M., Chidambaram, S.B., Guillemin, G.J., Sakharkar, M.K., 2020. Phosphodiesterase-4 enzyme as a therapeutic target in neurological disorders. Pharmacol. Res. 105078. https://doi.org/10.1016/j.phrs.2020.105078

Birben, E., Sahiner, U.M., Sackesen, C., Erzurum, S., Kalayci, O., 2012. Oxidative Stress and Antioxidant Defense. World Allergy Organ. J. 5, 9–19. https://doi.org/10.1097/WOX.0b013e3182439613

Boswell-Smith, V., Spina, D., Page, C.P., 2006. Phosphodiesterase inhibitors. Br. J. Pharmacol. 147, S252–S257. https://doi.org/10.1038/sj.bjp.0706495

Braidy, N., Grant, R., Adams, S., Brew, B.J., Guillemin, G.J., 2009a. Mechanism for quinolinic acid cytotoxicity in human astrocytes and neurons. Neurotox. Res. 16, 77–86. https://doi.org/10.1007/s12640-009-9051-z

Braidy, N., Grant, R., Adams, S., Brew, B.J., Guillemin, G.J., 2009b. Mechanism for quinolinic acid cytotoxicity in human astrocytes and neurons. Neurotox. Res. 16, 77–86. https://doi.org/10.1007/s12640-009-9051-z

Braidy, N., Grant, R., Adams, S., Guillemin, G.J., 2010. Neuroprotective effects of naturally occurring polyphenols on quinolinic acid-induced excitotoxicity in human neurons. FEBS J. 277, 368–382. https://doi.org/10.1111/j.1742-4658.2009.07487.x

Cao, X., Zhao, S., Liu, D., Wang, Z., Niu, L., Hou, L., Wang, C., 2011. ROS-Ca2+ is associated with mitochondria permeability transition pore involved in surfactin-induced MCF-7 cells apoptosis. Chem. Biol. Interact. 190, 16–27. https://doi.org/10.1016/j.cbi.2011.01.010

Cardinale, A., Fusco, F.R., 2018. Inhibition of phosphodiesterases as a strategy to achieve neuroprotection in Huntington’s disease. CNS Neurosci. Ther. 24, 319–328. https://doi.org/10.1111/cns.12834

Chandra, R., Aneja, R., Rewal, C., Konduri, R., Dass, S.K., Agarwal, S., 2000. An opium alkaloid-papaverine ameliorates ethanol-induced hepatotoxicity: Diminution of oxidative stress. Indian J. Clin. Biochem. 15, 155–160. https://doi.org/10.1007/BF02883745

Chen, Y., Stankovic, R., Cullen, K.M., Meininger, V., Garner, B., Coggan, S., Grant, R., Brew, B.J., Guillemin, G.J., 2010. The kynurenine pathway and inflammation in amyotrophic lateral sclerosis. Neurotox. Res. 18, 132–142. https://doi.org/10.1007/s12640-009-9129-7

Counts, S.E., Nadeem, M., Lad, S.P., Wuu, J., Mufson, E.J., 2006. Differential expression of synaptic proteins in the frontal and temporal cortex of elderly subjects with mild cognitive impairment. J. Neuropathol. Exp. Neurol. 65, 592–601. https://doi.org/10.1097/00005072-200606000-00007

D’Amelio, M., Cavallucci, V., Cecconi, F., 2010. Neuronal caspase-3 signaling: not only cell death. Cell Death Differ. 17, 1104–1114. https://doi.org/10.1038/cdd.2009.180

Dang, Y., Mu, Y., Wang, K., Xu, K., Yang, J., Zhu, Y., Luo, B., 2016. Papaverine inhibits lipopolysaccharide-induced microglial activation by suppressing NF-κB signaling pathway. Drug Des. Devel. Ther. 10, 851–859. https://doi.org/10.2147/DDDT.S97380

Dos Santos, A.A., López-Granero, C., Farina, M., Rocha, J.B.T., Bowman, A.B., Aschner, M., 2018. Oxidative stress, caspase-3 activation and cleavage of ROCK-1 play an essential role in MeHg-induced cell death in primary astroglial cells. Food Chem. Toxicol. Int. J. Publ. Br. Ind. Biol. Res. Assoc. 113, 328–336. https://doi.org/10.1016/j.fct.2018.01.057

Fernandez-Marcos, P.J., Auwerx, J., 2011. Regulation of PGC-1α, a nodal regulator of mitochondrial biogenesis1234. Am. J. Clin. Nutr. 93, 884S–890S. https://doi.org/10.3945/ajcn.110.001917

García, A.M., Redondo, M., Martinez, A., Gil, C., 2014. Phosphodiesterase 10 inhibitors: new disease modifying drugs for Parkinson’s disease? Curr. Med. Chem. 21, 1171–1187. https://doi.org/10.2174/0929867321666131228221749

Giampà, C., Laurenti, D., Anzilotti, S., Bernardi, G., Menniti, F.S., Fusco, F.R., 2010. Inhibition of the Striatal Specific Phosphodiesterase PDE10A Ameliorates Striatal and Cortical Pathology in R6/2 Mouse Model of Huntington’s Disease. PLOS ONE 5, e13417. https://doi.org/10.1371/journal.pone.0013417

Giralt, A., Saavedra, A., Carretón, O., Arumí, H., Tyebji, S., Alberch, J., Pérez-Navarro, E., 2013. PDE10 inhibition increases GluA1 and CREB phosphorylation and improves spatial and recognition memories in a Huntington’s disease mouse model. Hippocampus 23, 684–695. https://doi.org/10.1002/hipo.22128

Guillemin, G.J., 2012. Quinolinic acid, the inescapable neurotoxin. FEBS J. 279, 1356–1365. https://doi.org/10.1111/j.1742-4658.2012.08485.x

Guillemin, G. J., Brew, B.J., Noonan, C.E., Takikawa, O., Cullen, K.M., 2005. Indoleamine 2,3 dioxygenase and quinolinic acid immunoreactivity in Alzheimer’s disease hippocampus. Neuropathol. Appl. Neurobiol. 31, 395–404. https://doi.org/10.1111/j.1365-2990.2005.00655.x

Guillemin, G.J., Cullen, K.M., Lim, C.K., Smythe, G.A., Garner, B., Kapoor, V., Takikawa, O., Brew, B.J., 2007. Characterization of the kynurenine pathway in human neurons. J. Neurosci. Off. J. Soc. Neurosci. 27, 12884–12892. https://doi.org/10.1523/JNEUROSCI.4101-07.2007

Guillemin, Gilles J, Wang, L., Brew, B.J., 2005. Quinolinic acid selectively induces apoptosis of human astrocytes: potential role in AIDS dementia complex. J. Neuroinflammation 2, 16. https://doi.org/10.1186/1742-2094-2-16

Han, X., Lamshöft, M., Grobe, N., Ren, X., Fist, A.J., Kutchan, T.M., Spiteller, M., Zenk, M.H., 2010. The biosynthesis of papaverine proceeds via (S)-reticuline. Phytochemistry 71, 1305–1312. https://doi.org/10.1016/j.phytochem.2010.04.022

Hankir, M.K., Kranz, M., Gnad, T., Weiner, J., Wagner, S., Deuther□Conrad, W., Bronisch, F., Steinhoff, K., Luthardt, J., Klöting, N., Hesse, S., Seibyl, J.P., Sabri, O., Heiker, J.T., Blüher, M., Pfeifer, A., Brust, P., Fenske, W.K., 2016. A novel thermoregulatory role for PDE10A in mouse and human adipocytes. EMBO Mol. Med. 8, 796–812. https://doi.org/10.15252/emmm.201506085

Harada, A., Suzuki, K., Kimura, H., 2017. TAK-063, a Novel Phosphodiesterase 10A Inhibitor, Protects from Striatal Neurodegeneration and Ameliorates Behavioral Deficits in the R6/2 Mouse Model of Huntington’s Disease. J. Pharmacol. Exp. Ther. 360, 75–83. https://doi.org/10.1124/jpet.116.237388

Heckman, P.R.A., van Duinen, M.A., Bollen, E.P.P., Nishi, A., Wennogle, L.P., Blokland, A., Prickaerts, J., 2016. Phosphodiesterase Inhibition and Regulation of Dopaminergic Frontal and Striatal Functioning: Clinical Implications. Int. J. Neuropsychopharmacol. 19. https://doi.org/10.1093/ijnp/pyw030

Huttenlocher, P.R., Dabholkar, A.S., 1997. Regional differences in synaptogenesis in human cerebral cortex. J. Comp. Neurol. 387, 167–178. https://doi.org/10.1002/(sici)1096-9861(19971020)387:2<167::aid-cne1>3.0.co;2-z

Islam, M.T., 2017. Oxidative stress and mitochondrial dysfunction-linked neurodegenerative disorders. Neurol. Res. 39, 73–82. https://doi.org/10.1080/01616412.2016.1251711

Jabaris, S.G.S.L., Sumathy, H., Kumar, R.S., Narayanan, S., Thanikachalam, S., Babu, C.S., 2015. Effects of rolipram and roflumilast, phosphodiesterase-4 inhibitors, on hypertension-induced defects in memory function in rats. Eur. J. Pharmacol. 746, 138–147. https://doi.org/10.1016/j.ejphar.2014.10.039

Jourdi, H., Kabbaj, M., 2013. Acute BDNF Treatment Upregulates GluR1-SAP97 and GluR2-GRIP1 Interactions: Implications for Sustained AMPA Receptor Expression. PLOS ONE 8, e57124. https://doi.org/10.1371/journal.pone.0057124

Jung, J.-S., Shin, J.A., Park, E.-M., Lee, J.-E., Kang, Y.-S., Min, S.-W., Kim, D.-H., Hyun, J.-W., Shin, C.-Y., Kim, H.-S., 2010. Anti-inflammatory mechanism of ginsenoside Rh1 in lipopolysaccharide-stimulated microglia: critical role of the protein kinase A pathway and hemeoxygenase-1 expression. J. Neurochem. 115, 1668–1680. https://doi.org/10.1111/j.1471-4159.2010.07075.x

Kim, J.H., Yi, H.-J., Ko, Y., Kim, Y.-S., Kim, D.-W., Kim, J.-M., 2014. Effectiveness of papaverine cisternal irrigation for cerebral vasospasm after aneurysmal subarachnoid hemorrhage and measurement of biomarkers. Neurol. Sci. Off. J. Ital. Neurol. Soc. Ital. Soc. Clin. Neurophysiol. 35, 715–722. https://doi.org/10.1007/s10072-013-1589-0

Kim, S.M., Chung, M.J., Ha, T.J., Choi, H.N., Jang, S.J., Kim, S.O., Chun, M.H., Do, S.I., Choo, Y.K., Park, Y.I., 2012. Neuroprotective effects of black soybean anthocyanins via inactivation of ASK1-JNK/p38 pathways and mobilization of cellular sialic acids. Life Sci. 90, 874–882. https://doi.org/10.1016/j.lfs.2012.04.025

Kowiański, P., Lietzau, G., Czuba, E., Waśkow, M., Steliga, A., Moryś, J., 2018. BDNF: A Key Factor with Multipotent Impact on Brain Signaling and Synaptic Plasticity. Cell. Mol. Neurobiol. 38, 579–593. https://doi.org/10.1007/s10571-017-0510-4

Lee, Y.-Y., Park, J.-S., Leem, Y.-H., Park, J.-E., Kim, D.-Y., Choi, Y.-H., Park, E.-M., Kang, J.L., Kim, H.-S., 2019a. The phosphodiesterase 10 inhibitor papaverine exerts anti-inflammatory and neuroprotective effects via the PKA signaling pathway in neuroinflammation and Parkinson’s disease mouse models. J. Neuroinflammation 16, 246. https://doi.org/10.1186/s12974-019-1649-3

Lee, Y.-Y., Park, J.-S., Leem, Y.-H., Park, J.-E., Kim, D.-Y., Choi, Y.-H., Park, E.-M., Kang, J.L., Kim, H.-S., 2019b. The phosphodiesterase 10 inhibitor papaverine exerts anti-inflammatory and neuroprotective effects via the PKA signaling pathway in neuroinflammation and Parkinson’s disease mouse models. J. Neuroinflammation 16, 246. https://doi.org/10.1186/s12974-019-1649-3

Leenders, A.G.M., Sheng, Z.-H., 2005. Modulation of neurotransmitter release by the second messenger-activated protein kinases: Implications for presynaptic plasticity. Pharmacol. Ther. 105, 69–84. https://doi.org/10.1016/j.pharmthera.2004.10.012

Lim, C.K., Bilgin, A., Lovejoy, D.B., Tan, V., Bustamante, S., Taylor, B.V., Bessede, A., Brew, B.J., Guillemin, G.J., 2017. Kynurenine pathway metabolomics predicts and provides mechanistic insight into multiple sclerosis progression. Sci. Rep. 7, 1–9. https://doi.org/10.1038/srep41473

Miranda, M., Morici, J.F., Zanoni, M.B., Bekinschtein, P., 2019. Brain-Derived Neurotrophic Factor: A Key Molecule for Memory in the Healthy and the Pathological Brain. Front. Cell. Neurosci. 13. https://doi.org/10.3389/fncel.2019.00363

Mishra, J., Kumar, A., 2014. Improvement of mitochondrial function by paliperidone attenuates quinolinic acid-induced behavioural and neurochemical alterations in rats: implications in Huntington’s disease. Neurotox. Res. 26, 363–381. https://doi.org/10.1007/s12640-014-9469-9

Mizuno, T., Kurotani, T., Komatsu, Y., Kawanokuchi, J., Kato, H., Mitsuma, N., Suzumura, A., 2004. Neuroprotective role of phosphodiesterase inhibitor ibudilast on neuronal cell death induced by activated microglia. Neuropharmacology 46, 404–411. https://doi.org/10.1016/j.neuropharm.2003.09.009

Nazir, F.H., Becker, B., Brinkmalm, A., Höglund, K., Sandelius, Å., Bergström, P., Satir, T.M., Öhrfelt, A., Blennow, K., Agholme, L., Zetterberg, H., 2018. Expression and secretion of synaptic proteins during stem cell differentiation to cortical neurons. Neurochem. Int. 121, 38–49. https://doi.org/10.1016/j.neuint.2018.10.014

Niccolini, F., Haider, S., Reis Marques, T., Muhlert, N., Tziortzi, A.C., Searle, G.E., Natesan, S., Piccini, P., Kapur, S., Rabiner, E.A., Gunn, R.N., Tabrizi, S.J., Politis, M., 2015. Altered PDE10A expression detectable early before symptomatic onset in Huntington’s disease. Brain 138, 3016–3029. https://doi.org/10.1093/brain/awv214

Persson, J., Szalisznyó, K., Antoni, G., Wall, A., Fällmar, D., Zora, H., Bodén, R., 2020. Phosphodiesterase 10A levels are related to striatal function in schizophrenia: a combined positron emission tomography and functional magnetic resonance imaging study. Eur. Arch. Psychiatry Clin. Neurosci. 270, 451–459. https://doi.org/10.1007/s00406-019-01021-0

Radiske, A., Rossato, J.I., Gonzalez, M.C., Köhler, C.A., Bevilaqua, L.R., Cammarota, M., 2017. BDNF controls object recognition memory reconsolidation. Neurobiol. Learn. Mem., Memory reconsolidation and memory updating 142, 79–84. https://doi.org/10.1016/j.nlm.2017.02.018

Rahman, A., Rao, M.S., Khan, K.M., 2018. Intraventricular infusion of quinolinic acid impairs spatial learning and memory in young rats: a novel mechanism of lead-induced neurotoxicity. J. Neuroinflammation 15, 263. https://doi.org/10.1186/s12974-018-1306-2

Ren, J.-G., Seth, P., Everett, P., Clish, C.B., Sukhatme, V.P., 2010. Induction of Erythroid Differentiation in Human Erythroleukemia Cells by Depletion of Malic Enzyme 2. PLOS ONE 5, e12520. https://doi.org/10.1371/journal.pone.0012520

Rodefer, J.S., Saland, S.K., Eckrich, S.J., 2012. Selective phosphodiesterase inhibitors improve performance on the ED/ID cognitive task in rats. Neuropharmacology 62, 1182–1190. https://doi.org/10.1016/j.neuropharm.2011.08.008

Roush, E., Harlen, K., Hendrickson, M., Hughes, T.E., 2020. Neurodegenerative Disease and cAMP Signaling Dynamics. Biophys. J. 118, 456a. https://doi.org/10.1016/j.bpj.2019.11.2540

Schmidt, C.J., Chapin, D.S., Cianfrogna, J., Corman, M.L., Hajos, M., Harms, J.F., Hoffman, W.E., Lebel, L.A., McCarthy, S.A., Nelson, F.R., Proulx-LaFrance, C., Majchrzak, M.J., Ramirez, A.D., Schmidt, K., Seymour, P.A., Siuciak, J.A., Tingley, F.D., Williams, R.D., Verhoest, P.R., Menniti, F.S., 2008. Preclinical Characterization of Selective Phosphodiesterase 10A Inhibitors: A New Therapeutic Approach to the Treatment of Schizophrenia. J. Pharmacol. Exp. Ther. 325, 681–690. https://doi.org/10.1124/jpet.107.132910

Siuciak, J.A., McCarthy, S.A., Chapin, D.S., Fujiwara, R.A., James, L.C., Williams, R.D., Stock, J.L., McNeish, J.D., Strick, C.A., Menniti, F.S., Schmidt, C.J., 2006. Genetic deletion of the striatum-enriched phosphodiesterase PDE10A: evidence for altered striatal function. Neuropharmacology 51, 374–385. https://doi.org/10.1016/j.neuropharm.2006.01.012

Smith, S.M., Uslaner, J.M., Cox, C.D., Huszar, S.L., Cannon, C.E., Vardigan, J.D., Eddins, D., Toolan, D.M., Kandebo, M., Yao, L., Raheem, I.T., Schreier, J.D., Breslin, M.J., Coleman, P.J., Renger, J.J., 2013. The novel phosphodiesterase 10A inhibitor THPP-1 has antipsychotic-like effects in rat and improves cognition in rat and rhesus monkey. Neuropharmacology 64, 215–223. https://doi.org/10.1016/j.neuropharm.2012.06.013

Steffan, J.S., Kazantsev, A., Spasic-Boskovic, O., Greenwald, M., Zhu, Y.-Z., Gohler, H., Wanker, E.E., Bates, G.P., Housman, D.E., Thompson, L.M., 2000. The Huntington’s disease protein interacts with p53 and CREB-binding protein and represses transcription. Proc. Natl. Acad. Sci. 97, 6763–6768. https://doi.org/10.1073/pnas.100110097

Sumathi, T., Vedagiri, A., Ramachandran, S., Purushothaman, B., 2018. Quinolinic Acid-Induced Huntington Disease-Like Symptoms Mitigated by Potent Free Radical Scavenger Edaravone—a Pilot Study on Neurobehavioral, Biochemical, and Histological Approach in Male Wistar Rats. J. Mol. Neurosci. 66, 322–341. https://doi.org/10.1007/s12031-018-1168-1

Sundaram, G., Brew, B.J., Jones, S.P., Adams, S., Lim, C.K., Guillemin, G.J., 2014. Quinolinic acid toxicity on oligodendroglial cells: relevance for multiple sclerosis and therapeutic strategies. J. Neuroinflammation 11, 204. https://doi.org/10.1186/s12974-014-0204-5

Suzumura, A., Ito, A., Yoshikawa, M., Sawada, M., 1999. Ibudilast suppresses TNFalpha production by glial cells functioning mainly as type III phosphodiesterase inhibitor in the CNS. Brain Res. 837, 203–212. https://doi.org/10.1016/s0006-8993(99)01666-2

Valtschanoff, J.G., Burette, A., Davare, M.A., Leonard, A.S., Hell, J.W., Weinberg, R.J., 2000. SAP97 concentrates at the postsynaptic density in cerebral cortex. Eur. J. Neurosci. 12, 3605–3614. https://doi.org/10.1046/j.1460-9568.2000.00256.x

Wang, H., Xu, J., Lazarovici, P., Quirion, R., Zheng, W., 2018. cAMP Response Element-Binding Protein (CREB): A Possible Signaling Molecule Link in the Pathophysiology of Schizophrenia. Front. Mol. Neurosci. 11. https://doi.org/10.3389/fnmol.2018.00255

Wilson, R.F., White, C.W., 1986. Intracoronary papaverine: an ideal coronary vasodilator for studies of the coronary circulation in conscious humans. Circulation 73, 444–451. https://doi.org/10.1161/01.cir.73.3.444

Wu, Y., Chen, M., Jiang, J., 2019. Mitochondrial dysfunction in neurodegenerative diseases and drug targets via apoptotic signaling. Mitochondrion 49, 35–45. https://doi.org/10.1016/j.mito.2019.07.003

Yurtsever Kum, N., Yilmaz, Y.F., Gurgen, S.G., Kum, R.O., Ozcan, M., Unal, A., 2018. Effects of Parenteral Papaverine and Piracetam Administration on Cochlea Following Acoustic Trauma. Noise Health 20, 47–52. https://doi.org/10.4103/nah.NAH_31_17

Zinger, A., Barcia, C., Herrero, M.T., Guillemin, G.J., 2011. The Involvement of Neuroinflammation and Kynurenine Pathway in Parkinson’s Disease [WWW Document]. Park. Dis. https://doi.org/10.4061/2011/716859

